# ACE2 from *Pipistrellus abramus* bats is a receptor for HKU5 coronaviruses

**DOI:** 10.1101/2024.03.13.584892

**Authors:** Nicholas J. Catanzaro, Ziyan Wu, Chengcheng Fan, Alexandra Schäfer, Boyd L. Yount, Pamela J. Bjorkman, Ralph Baric, Michael Letko

## Abstract

The merbecovirus subgenus of coronaviruses includes Middle East Respiratory Syndrome Coronavirus (MERS-CoV), a zoonotic pathogen transmitted from dromedary camels to humans that causes severe respiratory disease. Viral discovery efforts have uncovered hundreds of merbecoviruses in different species across multiple continents, but few have been studied under laboratory conditions, leaving basic questions regarding their human threat potential unresolved. Viral entry into host cells is a critical step for transmission between hosts. Here, a scalable approach that assesses novel merbecovirus cell entry was developed and used to evaluate receptor use across the entire merbecovirus subgenus. Merbecoviruses are sorted into clades based on the receptor-binding domain of the spike glycoprotein. Receptor tropism is clade-specific, with the clade including MERS-CoV using DPP4 and multiple clades using ACE2, including HKU5 bat coronaviruses. Mutational analysis identified possible structural limitations to HKU5 adaptability and a cryo-EM structure of the HKU5-20s spike trimer revealed only ‘down’ RBDs.

## INTRODUCTION

Middle East Respiratory Syndrome Coronavirus (MERS-CoV) was first discovered in the middle east in 2012 as a cause of severe respiratory illness in humans with high mortality.^1^ Field studies have shown that MERS-CoV is endemic in dromedary camels in the region and routinely transmits to humans, sustaining a slow but continual regional outbreak.^2,3^ MERS-CoV is the prototypic member of the merbecovirus subgenus of beta-coronaviruses. Virus discovery efforts have uncovered hundreds of merbecoviruses circulating among diverse wildlife across multiple continents.^4–13^ As is true with most of the global virome, the majority of all sequenced coronaviruses have never been isolated or molecularly characterized and therefore their threat of spillover into humans remains unknown.

Coronaviruses must enter the cells and replicate to enable efficient transmission between species. MERS-CoV cell entry is mediated by the viral spike protein through several viral-host interactions: (1) the N-terminal domain of spike attaches virus to lectins present on the host cell surface, (2) the receptor binding domain (RBD) binds to the host cell receptor molecule, (3) and receptor binding reorganizes structural elements in the S2 subunit of the spike protein that mediate membrane fusion and subsequent cell entry.^14–16^ Each function is performed by a separate region of the spike protein, the RBD being responsible for the core interaction with the host receptor.^17–20^ While MERS-CoV uses dipeptidyl peptidase IV (DPP4) as its host receptor to infect cells, several studies have shown that some bat merbecoviruses do not use DPP4 and instead use angiotensin-converting enzyme 2 (ACE2) to infect their hosts.^12,21–24^

Similar to other beta-coronaviruses, the RBD in merbecovirus spike proteins is a single, contiguous domain located in the N-terminal half of the spike sequence, and it folds as a structurally stable unit that engages with the host receptor (Figure 1A-B).To reduce cost of viral gene synthesis while maintaining the ability to study viral protein function, we previously employed a method to replace the RBD of a sarbecovirus, SARS-CoV, with RBDs from other sarbecoviruses.^25^ The resulting chimeric spike gene was used in viral pseudotype assays to measure entry efficiency in cell culture of a panel of sarbecovirus RBDs.^25^ Using this approach (called “SarbecoType”), we previously characterized the cell entry for the majority of all published sarbecovirus RBDs, which led to isolation of new sarbecoviruses, identified understudied cell entry pathways, and helped elucidate the molecular determinants of sarbecovirus cell entry.^25–29^ We have also produced chimeric sarbecovirus molecular clones that are capable of replicating with a different receptor tropism when given either a different spike gene or RBD from other sarbecoviruses, further demonstrating the feasibility of exchanging the RBD within this subgenus.^30^

**Figure 1.**
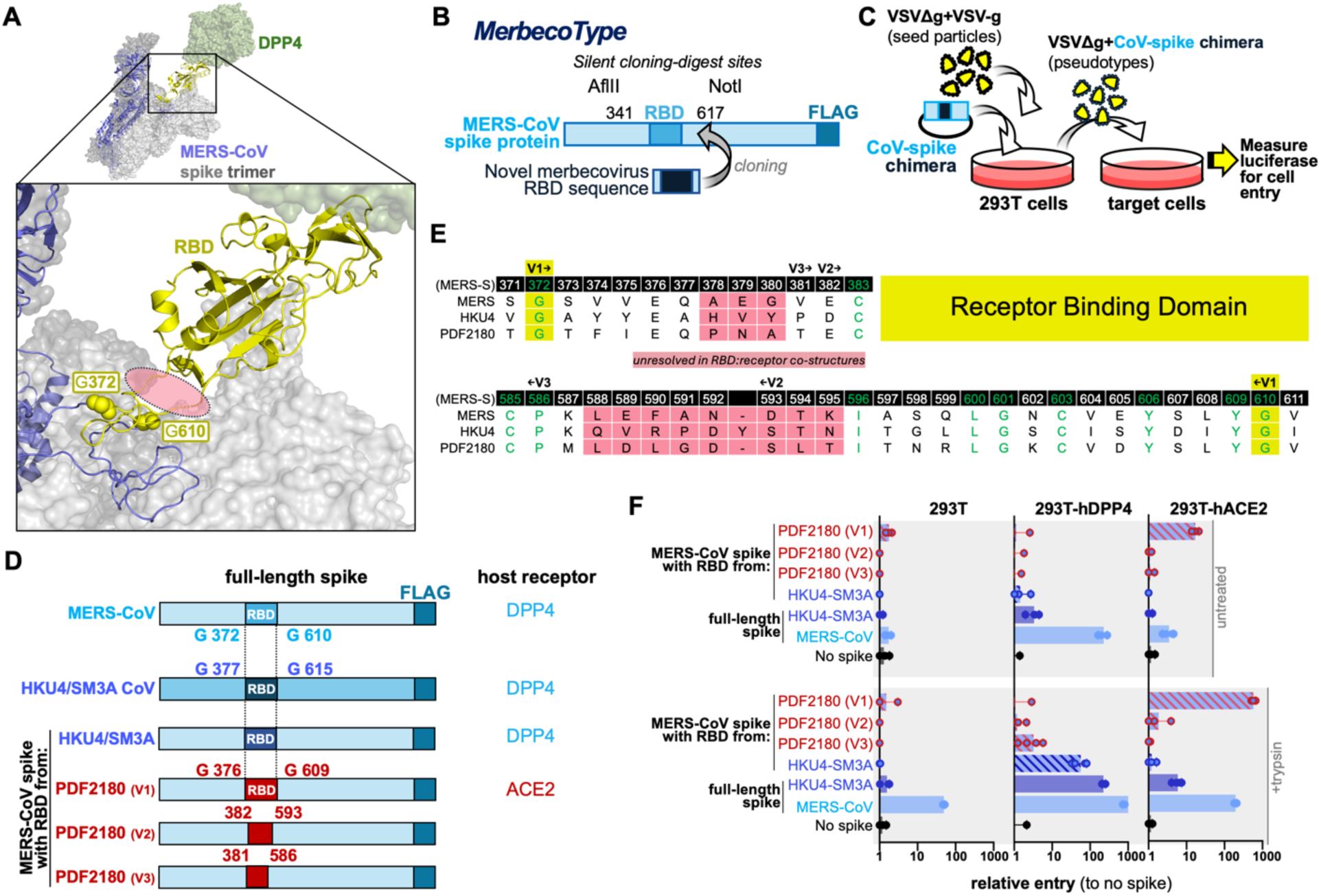
Identification of the functional RBD in merbecoviruses. **(A)** The receptor binding domain in merbecoviruses engages the host receptor, DPP4 (PDB ID: 4L72, 5X59). Conserved glycines marking the functional RBD boundaries are indicated in yellow spheres. **(B)** Overview of MERS-CoV spike design for RBD replacement. **(C)** Chimeric spikes are used to generate single-cycle pseudotyped VSV reporter particles. **(D)** Overview of chimeric spike designs. **(E)** Amino acid sequence alignment of region flanking the RBD in merbecoviruses. **(F)** 293T cell lines expressing indicated receptors were infected with pseudotypes and luciferase was measured for entry. Cells were infected in quadruplicate. For each graph, individual replicates are plotted as points, the mean is shown as a bar and lines indicate standard deviation.

Here, we took a similar approach, called “MerbecoType,” with the merbecovirus subgenus of betacoronaviruses. Similar to sarbecoviruses, RBDs of merbecoviruses can be classified into four different “clades” based on the presence or absence of indels, which exhibit clade-specific trends in receptor use and cell entry. Surprisingly, at least half of the merbecovirus RBD clades can use ACE2 for cell entry, with the entire group of HKU5 viruses exhibiting clear preference for ACE2 from their natural host species: *Pipistrellus abramus* bats and subsequently validated using live virus assays. We also determined the structure for HKU5 spike trimer and discuss similarities with other bat coronavirus spikes. These findings are important for ongoing efforts to identify looming coronavirus threats and in the development of interventions such as broad-spectrum antivirals and universal coronavirus vaccines.

## RESULTS

### Mapping the functional receptor binding domain

In order to test a large panel of RBDs from diverse merbecoviruses in viral pseudotype entry assays, MERS-CoV spike expression constructs were engineered to allow for easy replacement of the RBD with an RBD from other merbecoviruses (Figure 1A-C). MERS-CoV was selected as the recipient spike backbone because of its extensive structural characterization and compatibility with cell-culture based models of merbecovirus entry. ^1,31–33^ Similar to our prior sarbecovirus study,^25^ the MERS-CoV spike was codon optimized for human cells, appended with a C-terminal FLAG tag and silent mutations were introduced outside the RBD (Figure 1B). While MERS-CoV uses DPP4 for entry, ACE2 functions as a receptor for some other merbecoviruses including PDF2180 (Figure 1D).^22^ Thus the RBD from PDF2180 made an ideal candidate to develop a chimeric MERS-CoV spike and demonstrate successful receptor switching from DPP4 to ACE2 usage, generating a platform to explore different merbecovirus RBD receptor preferences in entry. To guide construction and expression of chimeric merbecovirus spikes, we utilized available structures. Unfortunately, amino acid motifs that are conserved across all merbecovirus spike sequences are rare, complicating the task of identifying common RBD exchange junctions across the merbecovirus subgenus. However, closer inspection of published spike sequences revealed several highly conserved glycine residues flanking the RBD (MERS-CoV spike positions 372 and 610; Figure 1A, D, E). These two residues flank the RBD region of spike used for structures with host receptor molecules.^22,34^ While the structures of RBDs have been determined for some merbecoviruses,^11,22,34,35^ regions flanking the RBD are flexible, making them difficult to structurally resolve in RBD:receptor complexes (Figure 1A, C; pink shaded region). To maximize chances for successful spike expression, several chimeras between MERS-CoV and PDF2180 were produced: version 1 included the region of PDF2180 spike between the conserved glycines, version 2 was based on structure and version 3 was based on the PDF2180-RBD as defined previously (Figure 1E).^22^ DNA fragments encoding the different RBD fragments were cloned into the MERS-CoV spike backbone and used to generate vesicular stomatitis virus (VSV) single-cycle pseudotypes (Figure 1C).

VSV-based pseudotypes were evaluated using a cell-based entry assay. While Baby Hamster Kidney cells (BHK cells) are not naturally susceptible to coronavirus entry but can support entry after ectopic receptor expression, BHK cells expressing human ACE2 do not always recapitulate natural viral entry for ACE2-dependent viruses with low affinity for the receptor.^29,36^ Therefore, 293T cells, which exhibit low susceptibility to coronavirus entry, were stably transduced to express human ACE2 or DPP4. Infecting these stable cell lines with pseudotypes bearing chimeric spikes revealed that the PDF2180 RBD was able to use human ACE2 for entry, indicating a successful receptor exchange for MERS-CoV spike (Figure 1D, F; PDF2180 V1). The regions of PDF2180 spike between RBD residues 376-381 and 586-609 were essential for receptor-use exchange, as two other spikes with shorter regions of the PDF2180 RBD were unable to use ACE2 (Figure 1A, D-F; PDF2180 V2, V3). Addition of exogenous protease (trypsin) during the assay enhanced viral entry, and afforded more sensitive detection of low affinity interactions. Importantly, trypsin-enhanced entry was still receptor dependent (Figure 1F). Production of a chimeric spike containing an analogous region from the DPP4-dependent HKU4-SM3A virus showed a retention in DPP4 use in MERS-CoV spike,^37^ further suggesting this region would allow for assessment of diverse receptor preferences (Figure 1E-F).

### Optimizing MERS-CoV spike for pseudotype assays

With an operational RBD-exchange strategy in place, the next goal was to optimize the spike backbone for pseudotype efficiency, which would allow for improved sensitivity to low-level viral entry. After engaging with the host receptor, coronavirus spikes are processed by host-cell proteases to release the spike fusion peptide and mediate membrane fusion.^38,39^ While the TMPRSS-family of cell-surface proteases and endosomal cathepsins have been shown to cleave many different coronavirus spikes, including MERS-CoV, SARS-CoV and SARS-CoV-2, furin is another protease that plays an important role in spike biogenesis and cell entry, depending on the availability of an appropriate cleavage site.^40–43^ MERS-CoV spike contains furin sites at the S1/S2 and S2’ junctions, although the role of furin in entry is less clear as removal of these sites has no effect on viral pseudotype entry *in vitro.*^38,44^ Because previous studies have shown that removal of the furin site from SARS-CoV-2 spike results in increased pseudotype efficiency,^45–47^ several versions of chimeric merbecovirus spikes were produced with either one or both furin sites mutated. Additionally, C-terminal truncations in the cytoplasmic tails of MERS-CoV, SARS-CoV and SARS-CoV-2 spikes increase surface expression of spike in transfected cells, resulting in improved pseudotype efficiency.^48–50^ Therefore, the protease site mutants were also tested with or without a 16-amino acid truncation in the C-terminal end of MERS-CoV spike (Figure S2A-C). Surprisingly, these modifications to spike had little effects on entry in 293T cells for MERS-CoV WT, PDF2180 RBD or HKU4/SM3A RBD (Figure S2B-C). While not statistically significant, spikes containing a C-terminal truncation and removal of the S2’ furin site performed slightly better than the other spikes; therefore, this set of modifications was included for further experiments.

### A pseudotype panel encompassing merbecovirus RBD diversity

The 2012 MERS-CoV outbreak renewed efforts in coronavirus sequencing from human patients and the dromedary camel intermediate host species, as well as virus discovery efforts. ^51–54^ Genbank has over 2000 entries for “merbecovirus” and “MERS-like” viruses; of these 2000 entries, but only about 842 entries contain full-length spike, and of those entries, approximately half are from species other than humans and camels (Figure 2A). Many coronavirus Genbank listings do not always contain species identifiers in the metadata, which left several MERS-CoV entries remaining even after excluding human and camel viruses. Further eliminating viruses titled as “MERS-CoV” from the list resulted in 65 spike sequences, which encompassed 34 unique RBD sequences (Figure 2A; Table 1, Figure S1). Some merbecoviruses are likely missing from this list as coronavirus nomenclature has evolved over the years. However, even with these considerations, this curated panel of merbecoviruses spans 18 years of virus discovery (some sequences in this study pre-date the MERS-CoV outbreak), 6 countries and 3 continents (Table 1).

**Figure 2.**
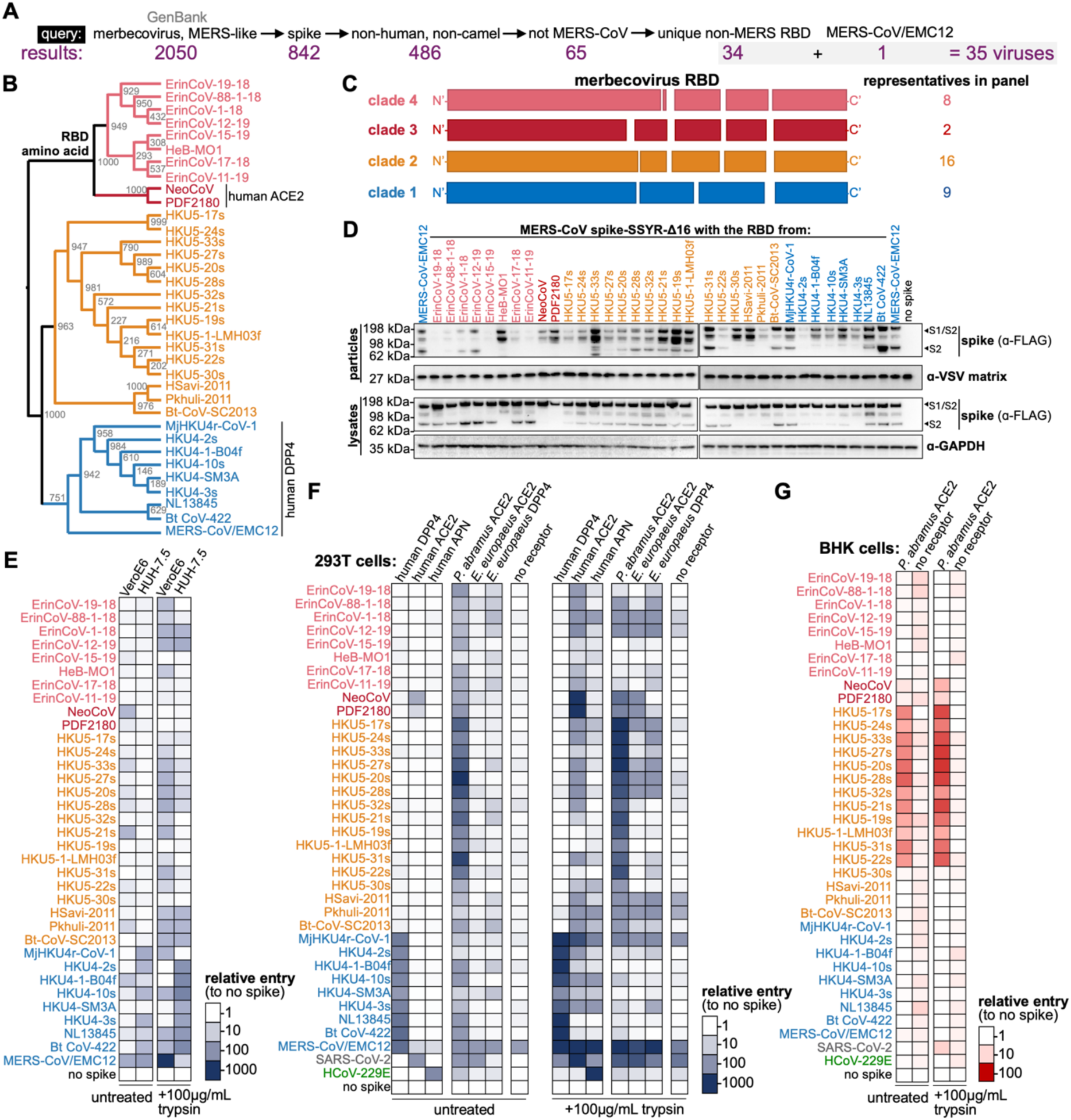
Merbecovirus RBDs sort into clades based on sequence and entry phenotype. **(A)** Total sequence diversity in merbecoviruses encompasses approximately 34 RBD sequences. **(B)** Cladogram showing the relationship of RBD amino acid sequences. Known receptors indicated. **(C)** Schematic overview of RBD clades, indicating the approximate size and location of indels characteristic to each merbecovirus RBD clade. **(D)** Western blot for FLAG (spike) expression of chimeric spikes. **(E)** VeroE6 cells or Huh-7.5 cells were infected with VSV pseudotype panel bearing indicated chimeric spikes, in quadruplicate. Luciferase was measured for cell entry. **(F)** 293T cells transduced to express human APN, human DPP4 or human ACE2 or transfected with *Erinaceus europaeus* ACE2, *Erinaceus europaeus* DPP4 or *Pipistrellus abramus* ACE2 were infected in the absence or presence of 100μg/mL trypsin. **(G)** BHK-21 cells were transiently transfected with empty vector or *Pipistrellus abramus* ACE2, and infected with or without exogenous trypsin. All cells were infected in quadruplicate. Heatmaps represent mean relative entry as compared to pseudotypes without spike.

**TABLE 1.** Diverse merbecovirus panel encompassing global diversity.

|  | Virus | Accession number | Receptor Clade | Host species | Location |
| --- | --- | --- | --- | --- | --- |
| 1 | ErinCoV-19-18 | MW246796 | 4 | *Erinaceus europaeus* | Italy |
| 2 | ErinCoV-88-1-18 | MW246795 | 4 | *Erinaceus europaeus* | Italy |
| 3 | ErinCoV-1-18 | MW246797 | 4 | *Erinaceus europaeus* | Italy |
| 4 | ErinCoV-12-19 | MW246800 | 4 | *Erinaceus europaeus* | Italy |
| 5 | ErinCoV-15-19 | MW246802 | 4 | *Erinaceus europaeus* | Italy |
| 6 | HKU31-HeB-MO1 | OM451213 | 4 | *Erinaceus amurensis* | China |
| 7 | ErinCoV-17-18 | MW246798 | 4 | *Erinaceus europaeus* | Italy |
| 8 | ErinCoV-11-19 | MW246799 | 4 | *Erinaceus europaeus* | Italy |
| 9 | NeoCoV | KC869678 | 3 | *Neoromicia capensis* | South Africa |
| 10 | PDF2180 | KX574227 | 3 | *Pipistrellus hesperidus* | Uganda |
| 11 | HKU5-17s | AGP04930 | 2 | *Pipistrellus abramus* | Hong Kong |
| 12 | HKU5-24s | AGP04936 | 2 | *Pipistrellus abramus* | Hong Kong |
| 13 | HKU5-33s | KC522104 | 2 | *Pipistrellus abramus* | Hong Kong |
| 14 | HKU5-27s | AGP04938 | 2 | *Pipistrellus abramus* | Hong Kong |
| 15 | HKU5-20s | AGP04933 | 2 | *Pipistrellus abramus* | Hong Kong |
| 16 | HKU5-28s | AGP04939 | 2 | *Pipistrellus abramus* | Hong Kong |
| 17 | HKU5-32s | AGP04942 | 2 | *Pipistrellus abramus* | Hong Kong |
| 18 | HKU5-21s | AGP04934 | 2 | *Pipistrellus abramus* | Hong Kong |
| 19 | HKU5-19s | AGP04932 | 2 | *Pipistrellus abramus* | Hong Kong |
| 20 | HKU5-1-LMH03f | NC009020 | 2 | *Pipistrellus abramus* | China |
| 21 | HKU5-31s | AGP04941 | 2 | *Pipistrellus abramus* | Hong Kong |
| 22 | HKU5-22s | AGP04935 | 2 | *Pipistrellus abramus* | Hong Kong |
| 23 | HKU5-30s | AGP04940 | 2 | *Pipistrellus abramus* | Hong Kong |
| 24 | HSavi-2011 | MG596802 | 2 | *Hypsugo savii* | Italy |
| 25 | PKhuli-2011 | MG596803 | 2 | *Pipistrellus kuhlii* | Italy |
| 26 | Bt-CoV-SC2013 | KJ473821 | 2 | *Vespertilio superans* | China |
| 27 | MjHKU4r-CoV-1 | OQ786862 | 1 | *Manis javanica* | China (Smuggled Malayan pangolins) |
| 28 | HKU4-2s | AGP04916 | 1 | *Tylonycteris pachypus* | Hong Kong |
| 29 | HKU4-1-B04f | NC009019 | 1 | *Tylonycteris pachypus* | Hong Kong |
| 30 | HKU4-10s | AGP04924 | 1 | *Tylonycteris pachypus* | Hong Kong |
| 31 | HKU4/SM3A | MW218395 | 1 | *Tylonycteris pachypus* | Hong Kong |
| 32 | HKU4-3s | AGP04917 | 1 | *Tylonycteris pachypus* | Hong Kong |
| 33 | NL13845 | MG021451 | 1 | *Tylonycteris pachypus* | China |
| 34 | Bt-CoV 422 | MG021452 | 1 | *Tylonycteris pachypus* | China |
| 35 | MERS/EMC12 | JX869059 | 1 | *Homo sapiens* | Saudi Arabia |

Phylogenetic analysis of the RBD amino acid sequences (as defined between the two glycine residues highlighted in Figure 1) has revealed several genetically-clustered groups based on characteristic indels and the sequences surrounding them (Figure 2B, C, Figure S1). The groups were labeled as clades 1 through 4: clade 1 includes MERS-CoV and HKU4 discovered between 2006-2012,^1,55,56^ clade 2 includes HKU5 also discovered in 2006,^55^ clade 3 incudes the African merbecoviruses, NeoCoV and PDF2180, discovered in 2017,^13^ and clade 4 includes the contemporary hedgehog merbecoviruses, “ErinCoVs,” discovered in Germany in 2014 and most recently identified in Italy in 2020 (Figure 2B-C, Table 1).^9,57–59^ RBDs for these merbecoviruses were synthesized and cloned into the optimized MERS-CoV spike backbone plasmid. All chimeric spike plasmids were stably expressed in 293T cells and used to produce pseudotyped viral-like particles for downstream experiments (Figure 2D). Concentrated supernatants were positive for the presence of spike by western blot analysis; however, variations observed in spike levels did not correlate with viral entry phenotypes observed in downstream assays (Figure 2D).

### Merbecovirus entry in common laboratory cell lines

Vero cells (from African Green Monkey kidneys) are common laboratory cell lines in coronavirus research and are used in attempts to isolate and propagate merbecoviruses and other viruses. ^1,60^ VeroE6 cells were poorly permissive for entry of the pseudotyped merbecovirus RBD panel, with addition of trypsin only mildly improving entry for some of the viruses in a calde-independent manner (Figure 2E). In contrast, Huh-7.5 cells, a human hepatocellular carcinoma cell line, were more susceptible to clade 1 merbecovirus RBDs (Figure 2E). Addition of trypsin improved entry for some of the clade 4 viruses for both cell lines (e.g., ErinCoV-1-18 and ErinCov-12-19) (Figure 2E).

### Merbecovirus compatibility with known human coronavirus receptors in human cells

The diverse panel of merbecovirus pseudotypes was tested on 293T cell lines stably transduced to express known human coronavirus receptors: amino peptidase N (APN), DPP4, or Angiotensin-ACE2. In the absence of trypsin, untransduced 293T cells exhibited poor susceptibility to the entire panel, while APN cells were susceptible to entry with a positive control: human coronavirus 229E, which is known to use APN as an entry receptor (Figure 2F).^61^ Cells transduced with human DPP4 exhibited clear susceptibility to clade 1 merbecovirus RBDs but not the other clades (Figure 2F), while cells expressing human ACE2 exhibited low level susceptibility to SARS-CoV-2 spike and clade 3 merbecovirus RBDs (Figure 2F). Exogenous trypsin moderately increased entry in untransduced 293T cells for human coronaviruses MERS-CoV and SARS-CoV-2, which uses human DPP4 and human ACE2, respectively, suggesting low levels of the human receptors were present on 293T cells (Figure 2F). In the presence of trypsin, clade 1 RBDs entered cells through DPP4, while clades 2, and 3 RBDs entered cells expressing ACE2 (Figure 2F). APN cells remained susceptible to HCoV-229E entry in the presence of trypsin. While entry signal under 10x is considered borderline permissive in this assay, and difficult to interpret, there were many viruses that presented low, but consistently detectable levels of ACE2 compatibility (Figure 2F).

### Species-specific receptor use across the merbecovirus RBD clades

Human cells expressing human ACE2 exhibited low level susceptibility to a wide range of merbeocviruses (Figure 2). To investigate if the merbecovirus RBD panel can use receptor orthologues from their cognate host species, ACE2 and DPP4 were synthesized from European hedgehogs (*Erinaceus europaeus*), the natural host species for the ErinCoVs (Table 1; merbecovirus RBD clade 4), and ACE2 was synthesized from Japanese house bats (*Pipistrellus abramus*): the natural host species for the HKU5 viruses (Table 1; merbecovirus RBD clade 2). DPP4 and ACE2 from European hedgehogs were inefficient as entry receptors and did not mediate entry for any of the merbecovirus clades in the absence of trypsin (Figure 2F). However, 293T cells transfected with ACE2 from Japanese house bats were highly permissive for entry by the entire HKU5 cluster of clade 2 merbecovirus RBDs – in the presence or absence of exogenous trypsin (Figure 2F).

To further reduce the background signal in the entry assays, 293T cells were exchanged with BHK cells, which are not susceptible to any sarbecovirus or merbecvirus due to significant hamster receptor incompatibilities and poor cell surface expression^62,63^. Following transfection with *Pipstrellus abramus* ACE2, BHK cells became highly permissive for HKU5 RBD-mediated entry (Figure 2G). Notably, all HKU5 sequences derive exclusively from *Pipistrellus abramus* samples (Table 1).

### Confirmation of HKU5 receptor use with stable cell lines and full-length spikes

The previous experiments were performed with human receptor stable cell lines and cells transiently transfected with *Pipistrellus abramus* ACE2. To achieve more consistent entry results with *Pipistrellus abramus* ACE2, a lentiviral vector was produced to generate stable cell lines, following the successful methodologies applied to generate the human receptor cell lines^29,64^ (Figure 3A). Unlike the parental cells, VeroE6 transduced with *Pipistrellus abramus* ACE2 were highly susceptible to the entire HKU5 RBD panel as well as the clade 3 viruses, NeoCoV and PDF2180, but remained poorly susceptible to clade 1, clade 4 and the clade 2 RBDs that were not identified in *Pipistrellus abramus* bats (compare Figure 2E compared with 3B). To further confirm receptor use for the HKU5 viruses, full-length spike sequences from HKU5-20s and HKU5-21s were synthesized and used to produce viral pseudotypes. For comparison, pseudotypes were also produced with the chimeric MERS-CoV-based spikes used in the previous experiments (Figure 3C). Full-length HKU5-20s and HKU5-21s spikes were expressed and incorporated into pseudotypes (Figure 3D). Again, while detection of spike varied across the concentrated pseudotype panel, it did not correspond with the observed entry results (Figure 3D-E). BHK cells transduced with *Pipistrellus abramus* ACE2 were susceptible to both full-length and chimeric HKU5-20s and -21s spikes but not MERS-CoV, a DPP4-dependent spike (Figure 3E, F). Altogether, the clear and species-specific entry observed for all HKU5 RBDs and full-length HKU5 spikes exclusively in cells expressing *Pipstrellus abramus* ACE2 strongly support a hypothesis that ACE2 is the natural receptor for these viruses in natural hosts.

**Figure 3.**
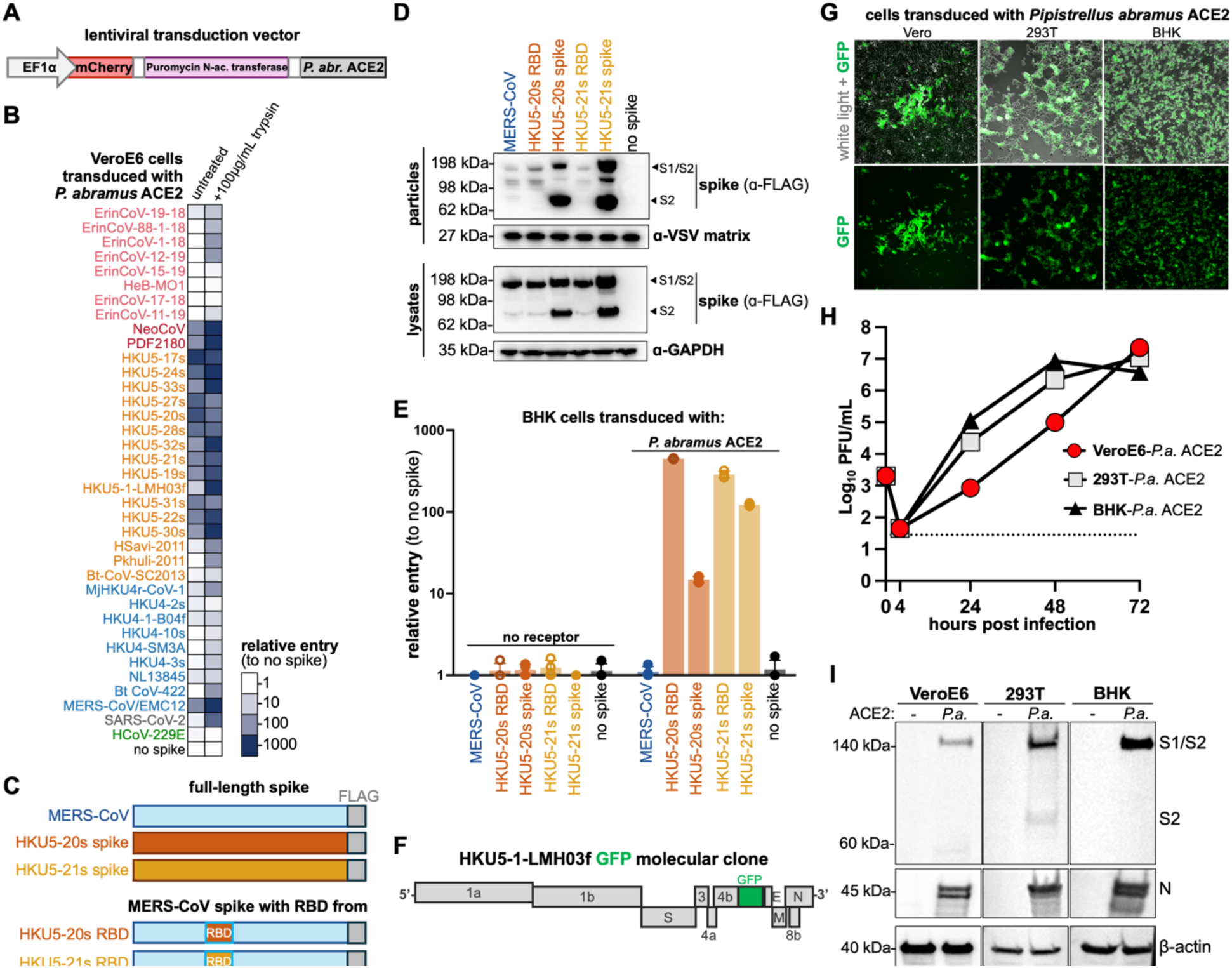
*Pipistrellus abramus* ACE2 rescues entry of full-length HKU5 spike in non-susceptible cells. **(A)** *Pipistrellus abramus* ACE2 expression cassette used in lentiviral transduction. **(B)** VeroE6 cells were transduced with *Pipistrellus abramus* ACE2 and infected with chimeric spike pseudotypes. **(C)** Schematic overview of full length and chimeric spikes. **(D)** Spike expression in cell lysates and incorporation in pseudotyped particles. **(E)** BHK-21 cells were transduced to express *Pipistrellus abramus* ACE2 and infected with pseudotypes bearing the indicated spikes. Cells were infected in quadruplicate. For each graph, individual replicates are plotted as points, the mean is shown as a bar and lines indicate standard deviation. **(F)** Overview of HKU5 molecular clone with GFP reporter in place of Orf5. **(G)** Light and fluorescent microscopy of indicated cell lines at 24-hours post-infection. **(H)** Growth curve of viral replication as measured by PFU of supernatants at indicated time points. **(I)** Western blot of infected cells for viral and host proteins at 24-hours post-infection.

### Confirmation of HKU5 receptor use with a replication-competent HKU5 molecular clone

To confirm if *Pipistrellus abramus* ACE2 could function as a receptor for true virus, we generated a replication competent molecular clone of HKU5 with green fluorescent protein (GFP) cloned in place of Orf5 (Figure 3F).^24^ Vero cells, 293T cells or BHK cells stably transduced with *Pipistrellus abramus* all strongly supported effecient viral replication in the absence of trypsin, as evidenced by GFP expression during replication, accumulation of infectious virus as measured by plaque assays, and production of viral spike and nucleocapsid proteins in infected cells (Figure 3G-I). Detection of distinct spike glycoprotein products at ∼180 and 75 kDa strongly support spike proteolytic processing and cleavage due to the presence of a putative furin cleavage site.

### Consensus sequences mimic clade-specific entry phenotypes

To further assess clade-specific receptor use, consensus sequences were generated for the RBDs from clades 1, 2 and 4 and used to produce chimeric spikes. PDF2180 was selected as a representative of clade 3 because clade 3 only contains two representative RBD sequences (Figure 4A). Pseudotypes bearing chimeric consensus RBD spikes generally reflected clade-specific entry phenotypes observed for individual RBDs from previous experiments (Figure 4B-C), suggesting that features common amino acids, structural features and motifs within each RBD clade dictate entry.

**Figure 4.**
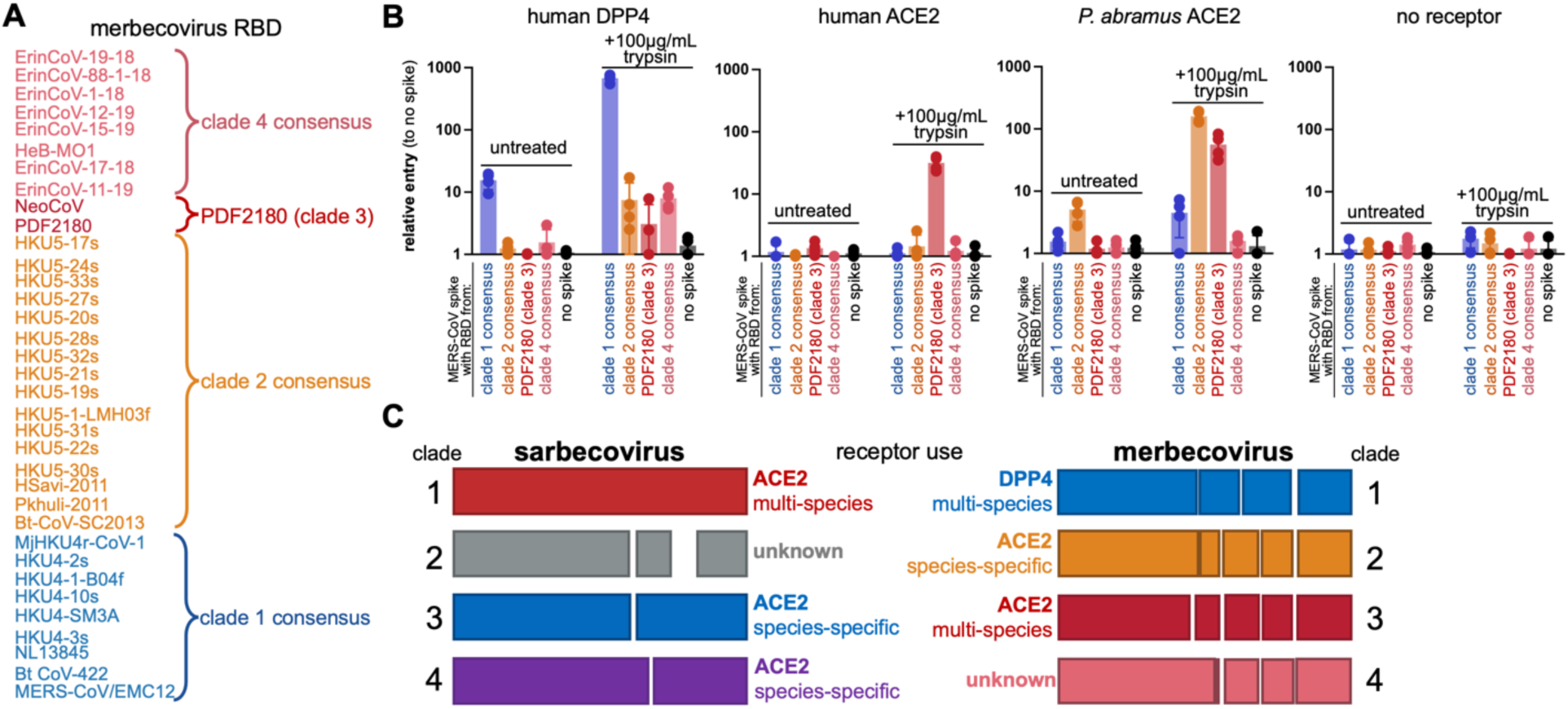
Clade consensus RBDs mimic clade-specific entry. **(A)** Consensus sequences were generated for clades 1, 2, and 4. **(B)** 293T cells transfected with indicated receptors were infected with chimeric spikes bearing clade consensus RBDs. **(C)** Schematic overview of betacoronavirus RBD clades, indel patterns and receptor use.

### Mutations in the HKU5 RBD modulate spike activity and species compatibility

A previous study reported structures of clade 3 merbecovirus RBDs from PDF2180 and NeoCoV bound to ACE2 from *Pipstrellus pipstrellus* bats.^22^ Although this study demonstrated HKU5 spike could not use *Pipistrellus pipistrellus* or any of 46 other bat ACE2s for entry, these RBD-ACE2 structures were used as a template to model predicted structures for HKU5 RBDs and *Pipistrellus abramus* ACE2 (Figure 5A-C). In the experimental structure, the interface between the clade 3 RBD, PDF2180, with *Pipistrellus pipistrellus* ACE2 involves multiple interactions with two surface-exposed loops on the RBD (Figure 5A), which we refer to here as Loop 1 and Loop 2. Curiously, the Loop 2 is highly polymorphic in merbecoviruses, with several HKU5 viruses, but not all, exhibiting a 4 amino acid deletion in this region (Figure 5B-C; Figure S1B). To assess if an analogous interface is compatible with interactions between HKU5 RBDs and bat ACE2, several sterically bulky or charged amino acid mutations were introduced at two points in the middle of Loops 1 and 2 (Figure 5). For these mutation experiments, HKU5-20s RBD was selected because it includes both loops and shows strong entry with *Pipistrellus abramus* ACE2, and HKU5-21s RBD was selected because it contains a 4 amino acid deletion in loop 2, while still strongly exhibiting entry using *Pipistrellus abramus* ACE2 (Figure 2, 3A, Figure S1C). For experimental consistency, BHK cells and 293T cells were transduced with a lentivector to constitutively express *Pipistrellus abramus* ACE2 and were infected in parallel assays to reduce effects from non-specific variation in the cell line background. Overall, mutations in Loop 2, but not Loop 1, of the HKU5 RBD disrupted entry with *Pipistrellus abramus* ACE2 (Figure 5E-G). While most of the mutations only reduced spike entry with bat ACE2, mutations in Loop 2 of the HKU5-20s RBD, but not the HKU5-21s RBD, increased entry with cells expressing human ACE2 (Figure 5G). While this difference was relatively small, the statistically-significant increase in entry was observed across different cell line backgrounds. Importantly, each mutation’s effect on viral entry was observed across both the BHK (hamster) and 293T-based (human) cell lines, suggesting the observed effects derive from the transduced ACE2 rather than other non-specific cell line variation (Figure 5).

**Figure 5.**
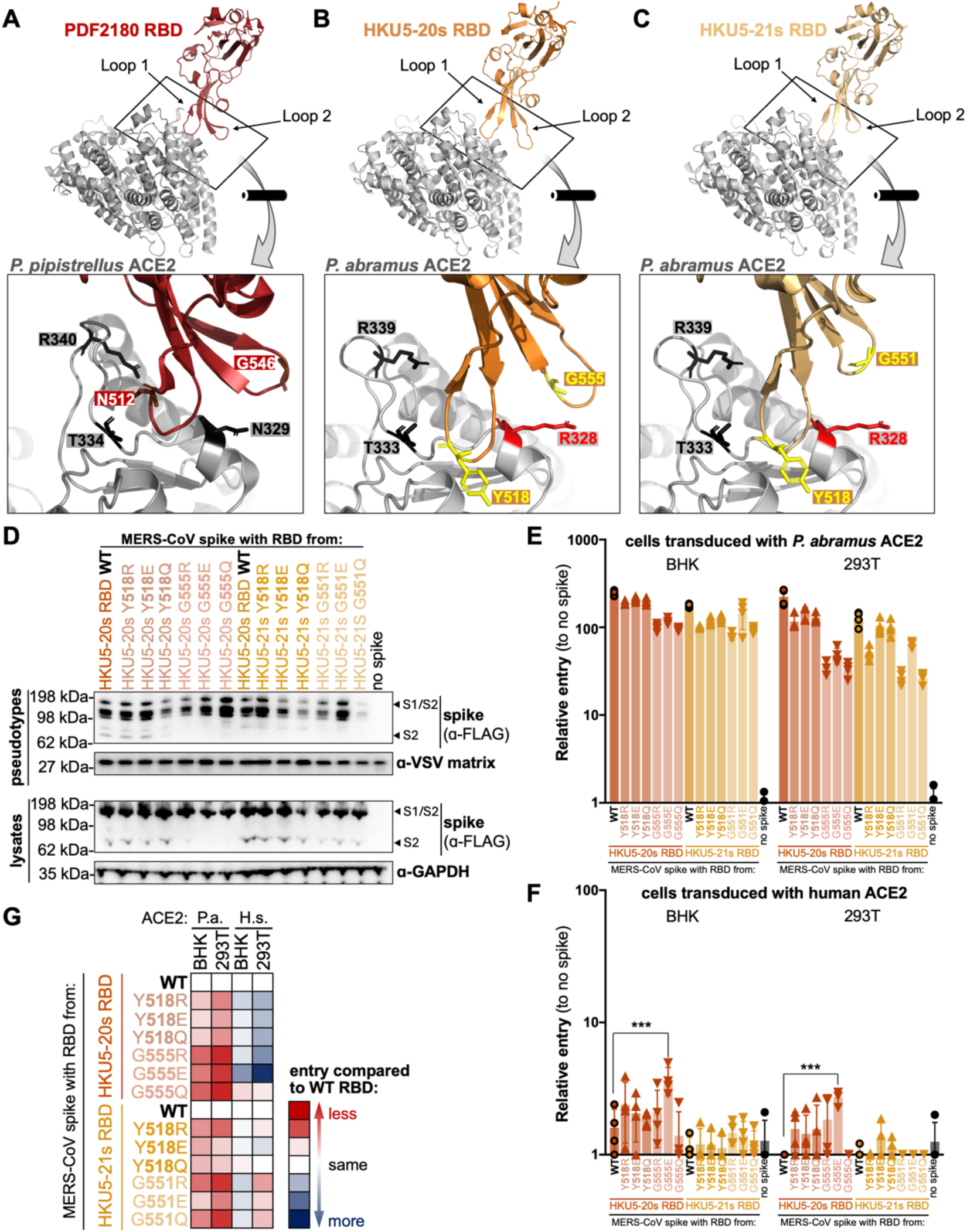
Structure-guided mutagenesis at the HKU5:ACE2 interface. **(A)** Co-structure for PDF2180 RBD and *Pipistrellus pipistrellus* ACE2 (PDB ID: 7wpz). **(B)** Predicted structures for HKU5-20s RBD and *Pipistrellus abramus* ACE2 (based on PDB IDs: 5xgr and 7c8j, respectively). **(C)** Predicted structure for HKU5-21s RBD and *Pipistrellus abramus* ACE2 (based on PDB ID: 8sak and 7c8j, respectively). **(D)** Mutant spike expression in cell lysates and incorporation in pseudotyped particles. **(E)** BHK or 293T cells transduced with either *Pipistrellus abramus* (P.a.) or **(F)** human (H.s.) ACE2 were infected with indicated pseudotypes. For each graph, four individual replicates are plotted as points, the mean is shown as a bar and lines indicate standard deviation. (***) Indicates P-value <0.001. **(G)** Entry values for mutant RBDs as compared to parental wild-type RBD.

### HKU5-20s spike structure reveals all ‘down’ RBDs

To further assess the interaction between HKU5-20s spike and *Pipistrellus abramus* ACE2 (paACE2), we attempted to solve a single-particle cryo-EM structure of a complex between a 2P stabilized^65^ KU5-20s trimeric ectodomain and soluble paACE-2 (both monomeric and dimeric paACE2-Fc), preparing grids at room temperature or at 37 °C. All grids yielded structures of the HKU5-20s spike trimer only, with no identifiable density for paACE2, suggesting further optimization of the binding conditions is need for determining a receptor-spike complex structure by cryo-EM. Indeed, prior work assessing merbecovirus spike interactions with *Pipistrellus* ACE2 were only in the context of purified RBD with receptor, not with a spike trimer ectodomain.^22^

We processed data from a grid incubated with paACE2-Fc plus HKU5-20s spike trimer ectodomain to derive a 2.8 Å structure of the unbound HKU5 trimer (Figure S3, Table S1). The structure revealed the expected S1 and S2 subunit organization as seen in other CoV spike trimers (Figure 6A-D). Although a distribution of ‘up’ and ‘down’ RBDs is usually found for sarbecovirus spike trimers,^66,67^ we found only ‘down’ RBDs in the HKU5-20s trimer structure (Figure 6A,B). To examine the degree of sequence conservation across 10 representative merbecovirus spikes (see Methods), we plotted amino acid sequence conservation on surface representations of structures of the HKU5-20s spike protomer and the HKU5-20s RBD (Figure 6D). We found more conserved regions in S2 and at the base of the RBD (Figure 6D). The well-resolved densities for the three ‘down’ RBDs allowed building of an RBD model including coordinates for Loops 1 and 2 (Figure 6E). These results provide additional structural insights for HKU5-20s spike trimer over an existing HKU4 RBD structure,^11^ and will be instrumental in future antigen design.

**Figure 6.**
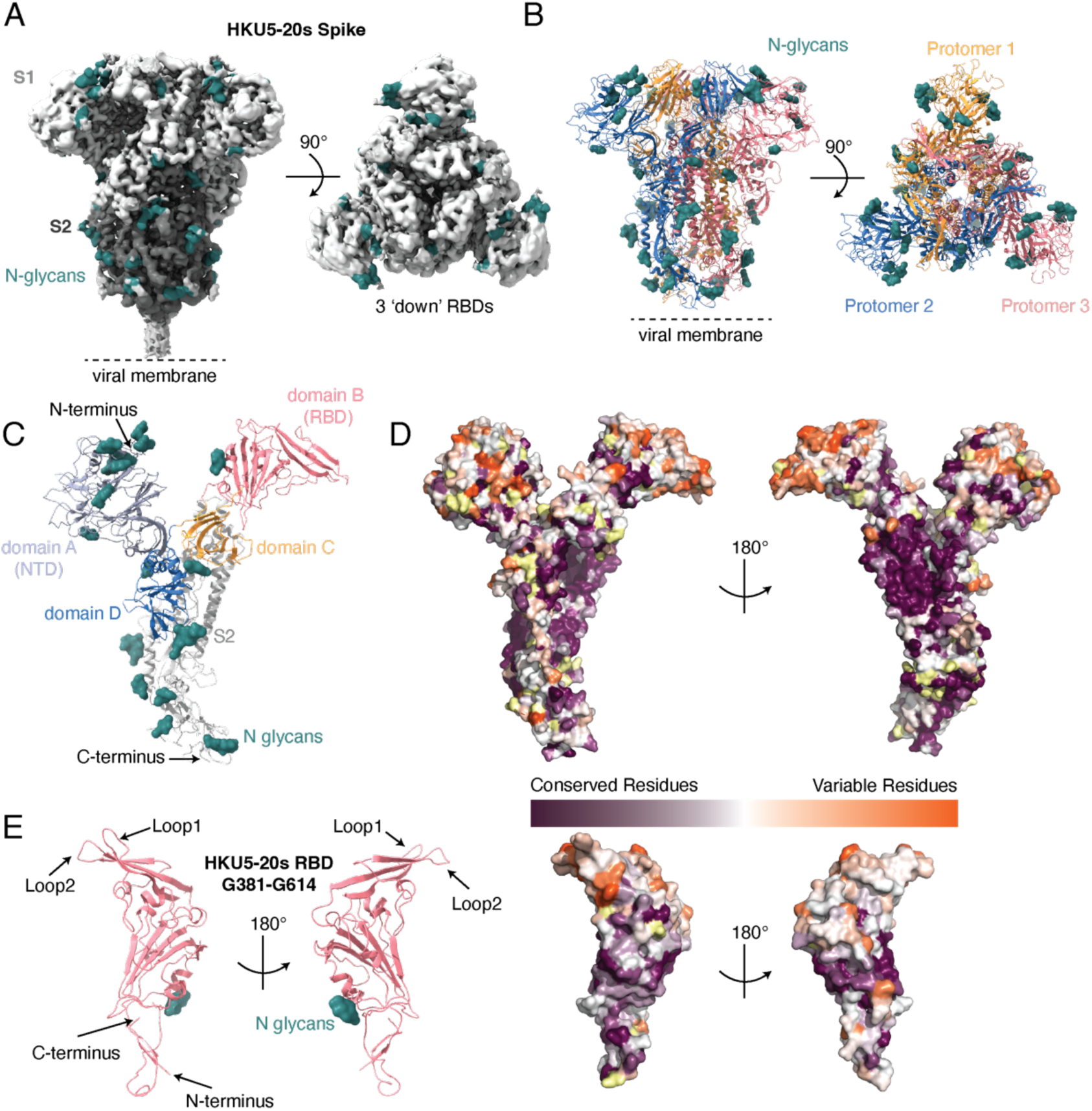
Structure of HKU5-20s spike trimer. **(A)** 2.8 Å EM density of structure of HKU5-20s spike with three ‘down’ RBDs is shown from the side (left) and top (right). **(B)** Cartoon diagram of HKU5-20s spike trimer with different colors for protomers shown from the side (left) and top (right). **(C)** Cartoon diagram of spike protomer showing domains A-D. **(D)** Amino acid sequence conservation of 10 merbecovirus spike sequences calculated by ConSurf Database^98^ plotted on a surface representation of an HKU5-20s protomer (top) and HKU5-20s RBD (bottom). Yellow patches in the purple to orange sequence conservation gradient indicate than <10% of the sequences were available for the calculation at that position. **(E)** Cartoon diagram of the HKU5-20s RBD from two orientations.

## DISCUSSION

Compared to sarbecoviruses, which share ≥90% nucleotide sequence identity in their RBDs,^25,68^ the diversity in merbecovirus RBD sequences is greater, initially complicating the RBD exchange strategy utilized in this study (Figure 1, 6E; Figure S1). Wide variability between merbecovirus host species (hedgehogs, bats, camels, humans and pangolins) also poses unforeseen challenges when studying cross-species interactions. The chimeric spike approach presented herein aims to eliminates some of the interspecies variability by using the human-compatible MERS-CoV spike as a backbone in experiments performed in human cells and other cells that support MERS-CoV infection. Our studies support the hypothesis that this approach reduces off-target phenotypes relating to cell attachment and protease processing. Addition of exogenous protease bypasses host-cell protease-processing, allowing for assessment of just the RBD interaction in the presence or absence of receptor. As now seen in this study and many others, chimeric coronavirus spikes faithfully re-create entry of both full spikes and wild-type viruses (Figure 1-3).^25–30,36,64,69^

Merbecovirus RBDs can be classified into four clades based on sequence, characteristic indels, and entry phenotype, but not geographic origin (Figure 2, 4; Figure S1, Table 1). Results from this study show that clade 1 viruses can use DPP4 from multiple species, clade 2 viruses have species-specific ACE2 preference, and clade 3 can use ACE2 from a wider range of host species. Recently, a merbecovirus found in Russian bats was shown to use ACE2 with a restricted host range, further demonstrating species-specific ACE2 interactions among merbecoviruses.^70^ No clear entry was observed with clade 4 viruses and receptors tested here, but nevertheless, human-cell entry was observed with ErinCoVs in the presence of trypsin (Figure 2E-G, 3B). Although the data presented here suggest ACE2 and DPP4 are not the receptor for clade 4 merbecoviruses, it is possible these viruses use these receptors in other host species not tested herein, and/or that the approach taken in this study is not suitable for detecting their entry. For example, if a region of ErinCoV spike outside of the RBD boundaries tested here interacts with the host receptor or regulates RBD binding, then the chimeric spike entry assays would miss this interaction. Additionally, while the chimeric spike strategy helps to reduce unforeseen entry barriers, dependence on extra-cellular factors remains possible, but is difficult to anticipate or control.^71^ Alternatively, clade 4 merbecovirus RBDs may use a receptor that is neither DPP4 nor ACE2, analogous to sarbecovirus RBD clade 2 viruses, which also can infect human cells through an unidentified route (Figure 2, 4).^25–28^

MERS-CoV spike can rapidly adapt to sub-optimal interactions with DPP4 from non-cognate species,^64^ and the ongoing SARS-CoV-2 pandemic underscores the ability of spike to adapt and correct sub-optimal ACE2 interactions while simultaneously escaping the immune response ^72–75^. Other research has shown that clade 3 merbecoviruses gain even broader host ACE2 tropism with a single point mutation in the spike RBD, T510F, significantly increasing tropism for human ACE2.^21,22^ Mutagenesis experiments presented here identified a region of HKU5 RBDs that also modifies ACE2 tropism (Figure 5). Structurally, T510F is on loop 1 of the RBD, while the effective mutations identified in this study are located on the adjacent loop 2 (Figure 5A-C). Only HKU5-20s, which possesses a larger second loop in its RBD, was able to gain consistent and statistically significant entry using human ACE2, with mutations. By contrast, HKU5-21s, with a shorter surface loop, did not measurably shift in ACE2 tropism (Figure 5E-G; Figure S1B). These results suggest that deletions within loop 2 of some HKU5 viruses may limit their evolutionary adaptability and that have a larger second loop, found in spikes such from viruses such as HKU5-20s, may allow for a wider range of ACE2 interactions. These data also suggest HKU5 spike may interact with a different region of ACE2 than has been reported for other merbecoviruses. Deep mutagenesis scanning and structural studies with both types of HKU5 RBDs are needed to further understand the HKU5-RBD interface, although even the low -level receptor compatibility reported here for these viruses is concerning and underscores the need for broadly protective coronavirus vaccines^69,76^.

Structural studies of other coronavirus spike trimers have revealed a distribution of different conformations with respect to the position of the RBDs.^11,67,77^ Coronavirus RBDs on a spike trimer can be in an ’up’ state, with receptor contacts being surface-exposed and capable of interacting with host cell receptors, or in the ‘down’ state, with the receptor contacts buried and inaccessible.^11,67,77^ MERS-CoV spike has been observed with one or two RBDs down,^78^ SARS-CoV spike has been observed with two or three ‘down’ RBDs,^78^ and SARS-CoV-2 spike has been observed with two or three ‘down’ RBDs.^66,67,79,80^ Analogously, the spike trimer from HCoV-229E, an alphacoronavirus with a tri-partite RBD, has been observed with all ‘down’ RBDs or some ‘up’ RBDs.^77^ Because the RBD is the primary target of neutralizing antibodies, transitions between ’up’ and ‘down’ RBD states is believed to play a role in immune evasion.^67,78^ We only observed HKU5-20s spike ectodomains with all ‘down’ RBDs (Figure 6). Structures of spike trimers from bat merbecoviruses PDF2180 and NeoCoV, as well as well as the bat sarbecovirus RsSHC014, have also been observed exclusively with all ‘down’ RBDs.^22,81^ Unlike the human respiratory coronaviruses, all bat coronaviruses have been identified in fecal or other gut-derived samples and are presumed to be gastrointestinal in their hosts.^6,12,13^ The low pH in the stomach and protease-rich gastrointestinal tract presents a unique and harsh environment for viruses, which may select for more closed confirmations in coronavirus spikes that have this tissue tropism. How these closed spikes transition to a state that can interact with the receptor is yet to be determined but may involve extracellular factors and/or pH changes.^66^

HKU5 viruses were first sequenced in 2006, shown to be DPP4-independent in 2014, and demonstrated to replicate in Vero cells in the presence of exogenous trypsin in 2020.^12,24,55^ To date, more than 40 ACE2 orthologues from bats and other species have been tested with HKU5 viruses;^6,22,24,82^ however, only ACE2 from *Pipistrellus abramus* has proven to mediate entry for these viruses (Figure 2-5). The RBD from HKU5 viruses shares ∼50% amino acid identity with MERS-CoV and clade 1 RBDs, while the RBD from NeoCoV is only ∼30-40% identical to MERS-CoV and HKU5 coronaviruses (Figure S1). This low sequence identity between NeoCoV and HKU5 RBDs has confounded efforts to predict the receptor for HKU5 from sequences alone.

A previous study identified ACE2 as the receptor for the clade 3 merbecoviruses such as PDF2180 and closely-related NeoCoV but was unable to identify the receptor for the clade 2 HKU5 viruses.^22^ This earlier study assessed the ability of HKU5 to interact with ACE2 from *Pipistrellus pipistrellus* bats and several other Yangochiropteran bats, but not the true host species for HKU5. The clear preference of HKU5 for *Pipistrellus abramus* ACE2, but not other ACE2 species orthologues tested clearly demonstrates a tighter species specificity found within clade 2 RBDs (Figure 2-3). *Pipistrellus abramus* ACE2 varies from previously tested *P. pipistrellus* and *P. khuli* ACE2s at residues 328 and 329, in close proximity to the interaction site previously reported with NeoCoV/PDF2180 RBDs, potentially explaining why HKU5 RBDs do not bind this orthologue (Figure 5A-C).^22^ Residue N329 in *Pipistrellus pipistrellus* ACE2 is also N-glycosylated,^21^ while the corresponding residue in *Pipistrellus abramus* ACE2, R328, is not, which may also explain the striking restriction in receptor use. Similar to the merbecovirus RBD clades, clade 1 sarbecoviruses can use ACE2 from diverse species, while clade 3 sarbecovirus RBDs only interact with ACE2 from their cognate species (Figure 4C).^36,69^

A clade 1 RBD pangolin merbecovirus has been shown to use human DPP4 and was isolated in Vero81 cells.^83^ While Vero cells are frequently used in virus isolation attempts,^1,60,84^ data from our study suggests that Huh-7.5 cells may be a more suitable cell line for isolating DPP4-dependent merbecoviruses and perhaps even some clade 4 merbecoviruses. The mechanisms underlying why these cells seem to allow more of these viruses to enter is yet to be determined. Furthermore, MERS-CoV grown in Vero cells has been shown to quickly accumulate tissue-culture adaptations in the spike protein driving the need for alternative cell culture models.^33^ Merbecoviruses represent yet another type of beta-coronavirus found in wildlife with potential human cell compatibility and a possibility to cause severe disease in humans. Japanese house bats (*Pipistrellus abramus*) are the natural host for HKU5 coronaviruses and are synanthropic, roosting in human dwellings and structures, and therefore pose repeat opportunities for human exposure.^55,85^ Luckily, clade 2 merbecovirus RBDs exhibit low human ACE2 compatibility (Figure 2-5), although efforts should still be made to reduce contact with these animals. The development of replication competent HKU5 and PDF2180 recombinant viruses will support the development of pan-coronavirus and pan-merbecovirus vaccines and small molecule countermeasures to protect the health of populations globally. The work presented here will serve as a guide to understanding merbecovirus cell entry, subsequent spillover risk and support One Health Preparedness.

## METHODS

### Biosafety

Prior approval for experiments using full-length, recombinantly derived HKU5 was obtained from the Institutional Biosafety Committee of the University of North Carolina at Chapel Hill. All experiments using live HKU5 virus were performed under Biosafety Level 3 conditions with personnel wearing full-body personal protective equipment and HEPA-filtered respiratory protection.

### Cells and viruses

293T, VeroE6, Huh-7.5 and BHK-21 cells were maintained in DMEM supplemented with 10% FBS, penicillin-streptomycin, and L-glutamine and maintained at 37°C, 5% CO2. Cells stably expressing coronavirus receptors were generated as previously described and maintained under selection with either 1μg/mL puromycin (BHK and 293T) or 5μg/mL puromycin (VeroE6).^29^ All cell lines were species-confirmed with cytochrome sequencing and verified mycoplasma negative with MycoSniff PCR Kit (MP Bio). The molecular clone derived HKU5-GFP was generated as previously described.^24^ Briefly, genomic cDNA sequences were ligated, *in vitro* transcribed, and electroporated into BHK cells. Stocks of recombinant HKU5-GFP were then propagated in Vero cells stably expressing *Pipistrellus abramus* ACE2 at 37°C and 5.0% CO2. For multistep growth curve analysis, cells were infected at an MOI of 0.01. Samples of infected cell culture supernatant were collected at 0, 4, 24, 48, and 72 h.p.i. and stored at -80 for titration via plaque assay. For fluorescent microscopy and western blot experiments, cells were infected with HKU5-GFP at an MOI of 1.0. At 24 h.p.i., GFP-fluorescence was monitored using an inverted optical microscope (Olympus, IX73), and lysates were harvested for western blot analysis as described below.

### Phylogenetic analysis

As described previously, the amino acid sequences for spike receptor binding domains aligned using Clustal multiple sequence alignment software and default parameters. A maximum likelihood phylogenetic tree was produced with PhyML v. 3.0 ^86^. The ‘WAG’ matrix +G model of amino acid substitution was selected by the Smart Model Selection method with 1000 bootstrap replicates^87^. A tree was then visualized as a cladogram with FigTree v1.4.4 (https://github.com/rambaut/figtree). Clade consensus sequences were generated with the aid of Geneious (Dotmatics, Inc.).

### Plasmids

MERS-CoV/EMC12 spike (accession number JX869059) was codon optimized, appended with C-terminal FLAG tag, and silent mutations were introduced near spike amino acids 341 and 617 to generate AflII and NotI restriction digest sites. RBDs were codon optimized and synthesized (IDT DNA) as previously described for sarbecoviruses^25^. Lyophilized RBD fragments were resuspended in water and used in downstream Infusion based cloning (Takara Bio) reactions to assemble full-length chimeric spike proteins. *Erinaceus europaeus* ACE2, transcript variant 1 (genbank XM_060183012.1), *Pipistrellus abramus* ACE2 (genbank GQ262782.1) and *Erinaceus europaeus* DPP4 (genbank XM_060177812.1) were codon optimized, split in half and synthesized for Infusion based DNA assembly. Splitting genes into multiple parts reduces synthesis time, costs and overall synthesized product complexity. A complete list of accession numbers for the spike panel can be found in Table 1.

### *Pipistrellus abramus* ACE2 cell lines

*Pipistrellus abramus* ACE2 was sub-cloned from expression plasmid into a lentiviral expression vector with PCR and used to produce lentiviral particles in 293T cells ^64,88^. Supernatants were filtered, aliquoted and frozen at -80°C. BHK cells, 293 T cells and VeroE6 cells were transduced with a lentiviral expression vector as previously described ^29,64^. Cells were maintained for 3 passages under selective pressure before use in experiments.

### Pseudotype entry assay

Pseudotype assays were performed as previously described^26,27^. For human ACE2, APN, and DPP4 experiments, 293T cells stably transduced with potential coronavirus receptors were seeded in black 96-well plates and subsequently infected 24 hours later with equal volumes of viral pseudotypes. Infections were performed on ice to prevent unintended, premature trypsin activity. Pseudotypes were combined with either HBSS or HBSS containing trypsin (not TPCK-treated) to achieve a final concentration of 100μg/mL trypsin. Cells were washed once with cold PBS, centrifuged with pseudotypes at 4°C, 1200 x g for 1 hour and incubated at 37°C overnight. Luciferase was measured using Bright-Glo reagent (Promega) approximately 18 hours post-infection. Relative entry was calculated by dividing raw luciferase values for each spike by the signal for no-spike pseudotypes. To compensate for plate reader background error, plates were measured 4 times, analyzed individually and then averaged across all four measurements. Relative entry values for all four replicates were averaged and plotted as heatmaps using Microsoft Excel.

### Statistics and reproducibility

Each figure shows data that is representative of four technical replicates. During this study, experiments were performed with different batches of pseudotypes and cells and at different times. Therefore, the results shown are representative of the observed biological replicates. Statistical significance for mutation comparisons in Figure 5 were determined by 2-way ANOVA with Tukey correction for multiple comparisons (GraphPad).

### Structure prediction and docking

MERS-CoV spike RBD bound to DPP4 (PDB ID: 4L72) ^35^, the MERS-CoV spike trimer (PDB ID: 5X59)^78^, and the co-structure of NeoCoV-RBD with P. pipistrellus ACE2 (PDB ID: 7WPZ)^22^, were visualized in PyMol (version 2.4.0). Structures for HKU5-20s and HKU5-21s RBDs were predicted with SwissModel using PDB IDs 8SAK and 5XGR, respectively^34,89^. *Pipistrellus abramus* ACE2 was modelled with similar methods using PDB ID 7C8J as a guide^90^.

### Protein Expression for cryo-EM

A trimeric HKU5-20s trimeric ectodomain (residues 24-1301 of the GenBank sequence AGP04933) with 2P stabilizing mutations,^65^ a mutated furin cleavage site between S1 and S2, a C-terminal TEV site, foldon trimerization motif, 8xHis tag, and AviTag) was expressed as described.^91^ A gene encoding a 8xHis-tagged soluble paACE2 (*Pipistrellus abramus* ACE2) construct (residues 20-613 of the Genbank sequence ACT66266) or a soluble paACE2-Fc (residues 20-613 of Genbank sequence ACT66266 fused to human IgG1 Fc) were expressed and purified as described for human ACE2 and human ACE2-Fc^92^ by nickel-NTA (ACE2) or protein A (paACE2-Fc) chromatography followed by size-exclusion chromatography using a Superdex 200 column (Cytiva) as described.^93^ Peak fractions were identified by SDS-PAGE, and those containing spike trimer, paACE2, or paACE2-Fc were pooled, concentrated, and stored at 4 °C until use.

### Cryo-EM sample preparation

paACE2-spike trimer and paACE2-Fc–spike trimer complexes were prepared by incubating purified paACE2 or paACE2-Fc with HKU5-20s trimer at a molar ratio of 3:1 (paACE2 or pACE2-Fc:spike) at 37°C for 30 minutes to a final concentration of ∼2 mg/mL. Cryo-EM grids were prepared using a Mark IV Vitrobot (ThermoFisher) at 37 °C and 100% humidity. Sample (3 μL) was applied to 300 mesh Quantifoil R1.2/1.3 grids (Electron Microscopy Sciences) that had been freshly glow discharged with PELCO easiGLOW (Ted Pella) for 1 min at 20 mA, blotted for 3 seconds with Whatman No.1 filter paper, and grids were vitrified in liquid ethane. Tweezers were preheated to 37°C before picking up grids.

### Cryo-EM data collection and processing

A single-particle cryo-EM dataset for HKU5-20s spike 2P with paACE2-Fc was collected using SerialEM automated data collection software^94^ on a 200 keV Talos Arctica cryo-electron microscope (Thermo Fisher Scientific) equipped with a K3 camera (Gatan) (Table S1). Movies were recorded with 40 frames, a defocus range of -1 to -3 μm, and a total dosage of 60 e-/Å^2^ using a 3x3 beam image shift pattern with 3 exposures per hole in the super-resolution mode with a pixel size of 0.4345 Å. The dataset was motion corrected with patch motion correction using a binning factor of 2, and CTF parameters were estimated using Patch CTF in cryoSPARC v4.5.3.^95^ Particle picking was done with blob picker in cryoSPARC using a particle diameter of 100 to 200 Å, and movies and picked particles were inspected before extraction. Particles were extracted and classified using 2D classification in cryoSPARC. Particles from selected classes were used for template picker with a particle diameter of 150 Å. Picked particles were inspected before extraction and 2D classification. After exposure curation, the remaining particles from selected classes were used for ab initio modeling with 4 volumes, which were further refined with heterogeneous refinement in cryoSPARC. Subsequent homogeneous and non-uniform refinements were carried out, and the final reconstruction was obtained from a non-uniform refinement conducted with C3 symmetry in cryoSPARC.

Structure figures were made using ChimeraX.^96^ For plotting sequence conservation onto protein structure surfaces, sequence alignments of 10 representative merbecovirus spikes were performed using Geneious Prime. Conservation scores for analogous residues (HKU5-20s residues 24-1231 for spike protomer and residues 381-614 for RBD) were calculated and mapped onto the HKU5-20s protomer or RBD by using ConSurf.^97^ The following 10 merbecovirus spike sequences were used in the sequence alignment:

- GenBank AFS88936 (MERS-CoV-EMC/2012)
- GenBank AGP04933 (HKU5-20s)
- GenBank AGP04934 (HKU5-21s)
- GenBank AGY29650 (NeoCoV)
- GenBank AHY61337 (BtVs-BetaCoV/SC2013)
- GenBank ARJ34226 (PDF2180)
- GenBank AUM60014 (Bat-CoV/H.savii)
- GenBank AUM60024 (Bat-CoV/P.khulii)
- GenBank QRN68031 (ErinCoV-19/2018)
- GenBank USL83011 (MOW15-22/2015)

MERS-CoV/EMC12 was selected because of high sequence identity with HKU5 spikes;^12^ NeoCoV, PDF2180 and MoW-15-22 spikes were selected because they have been shown to use ACE2;^21,22,70^ BatCoV/PKhuli, H.Savii, and SC2013 were selected as other clade 2 RBDs to compare with HKU5 RBD (Figure S1), and the clade 4 RBD from ErinCoV-19-18 was selected because of low sequence identity with other merbecovirus RBD clades (Figure S1).

### Western blot

As described previously,^25^ 293T cells transfected with spike plasmids were lysed in 1% SDS lysis buffer and stored at -80°C until use. Lysates were clarified, boiled, reduced and separated on 10% Bis-Tris gels in MOPS buffer (Invitrogen NuPage).^25^ Pseudotypes were concentrated over iodixanol (Opti-Prep; Sigma) gradients as previously described ^25,28,29^. Membranes were probed with FLAG:HRP conjugated monoclonal antibody (ThermoFisher), GAPDH monoclonal (ThermoFisher), MERS-CoV N (Sino Biological; catalog number: 40068-RP02; 1:2000), MERS-CoV S2 (Invitrogen; catalog number: 66008-1-Ig; 1:5000) and reprobed with a secondary goat-anti-mouse:HRP poloyclonal mix (ThermoFisher), goat-anti-rabbit-HRP (Cell Signaling Technology; catalog number: 7074S; 1:5000), or goat anti-mouse-HRP (Cell Signaling Technology; catalog number: 7076S; 1:5000).

## Data availability

Data that support the findings of this study are available from the corresponding author upon request. Accession numbers for all spike sequences used in this study can be found in Table 1. Atomic models and cryo-EM maps generated from cryo-EM studies of the HKU5-20s spike have been deposited at the Protein Data Bank (PDB) and Electron Microscopy Data Bank (EMDB) under accession codes PDB 9D4T and EMBD-46569, respectively

## Reagent availability

All biological reagents are available upon request.

## Acknowledgements

We would like to thank Dr. Stephanie Seifert, Dr. Alex Cohen, Dr. Tyler Starr and Dr. Thomas Gallagher for providing valuable insights and comments on the work and manuscript. Research reported in this publication was supported by the National Institute of Allergy and Infectious Diseases of the National Institutes of Health (NIAID/NIH) under Award Numbers R01AI179720 to M.L., R01 AI167966 and CEIRS Contract 75N93021C00014 to R.S.B., and P01AI100148 to P.J.B. The content is solely the responsibility of the author and does not necessarily represent the official views of the National Institutes of Health

---

**Figure S1.**
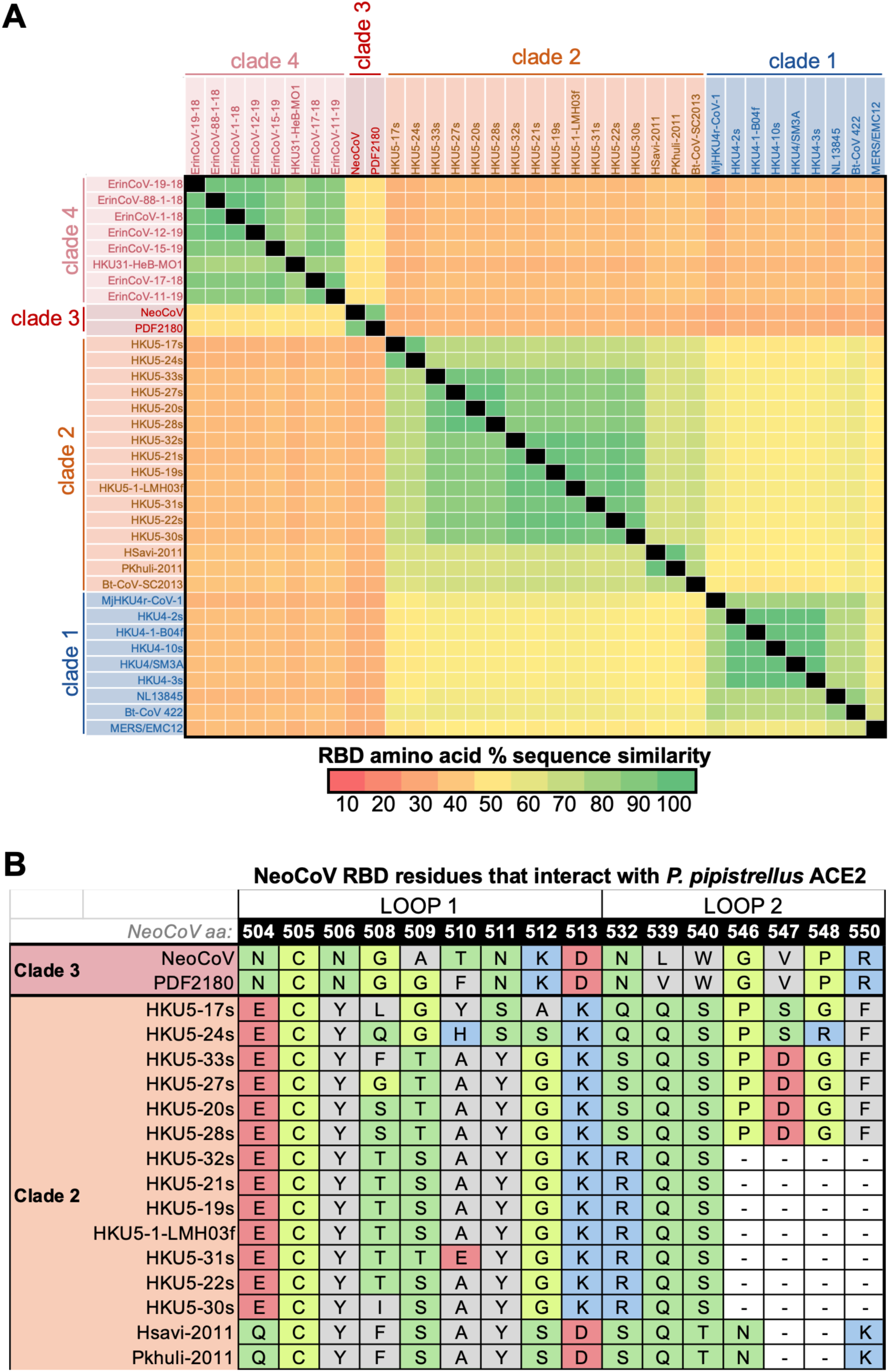
Sequence Identity matrix of merbecovirus RBDs. **(A)** Amino acid sequences for each spike RBD were aligned and compared. Black squares indicate identical residue. **(B)** Comparison of clade 3 RBD residues that interact with *P. pipistrellus* ACE2 with clade 2 sequences.

**Figure S2.**
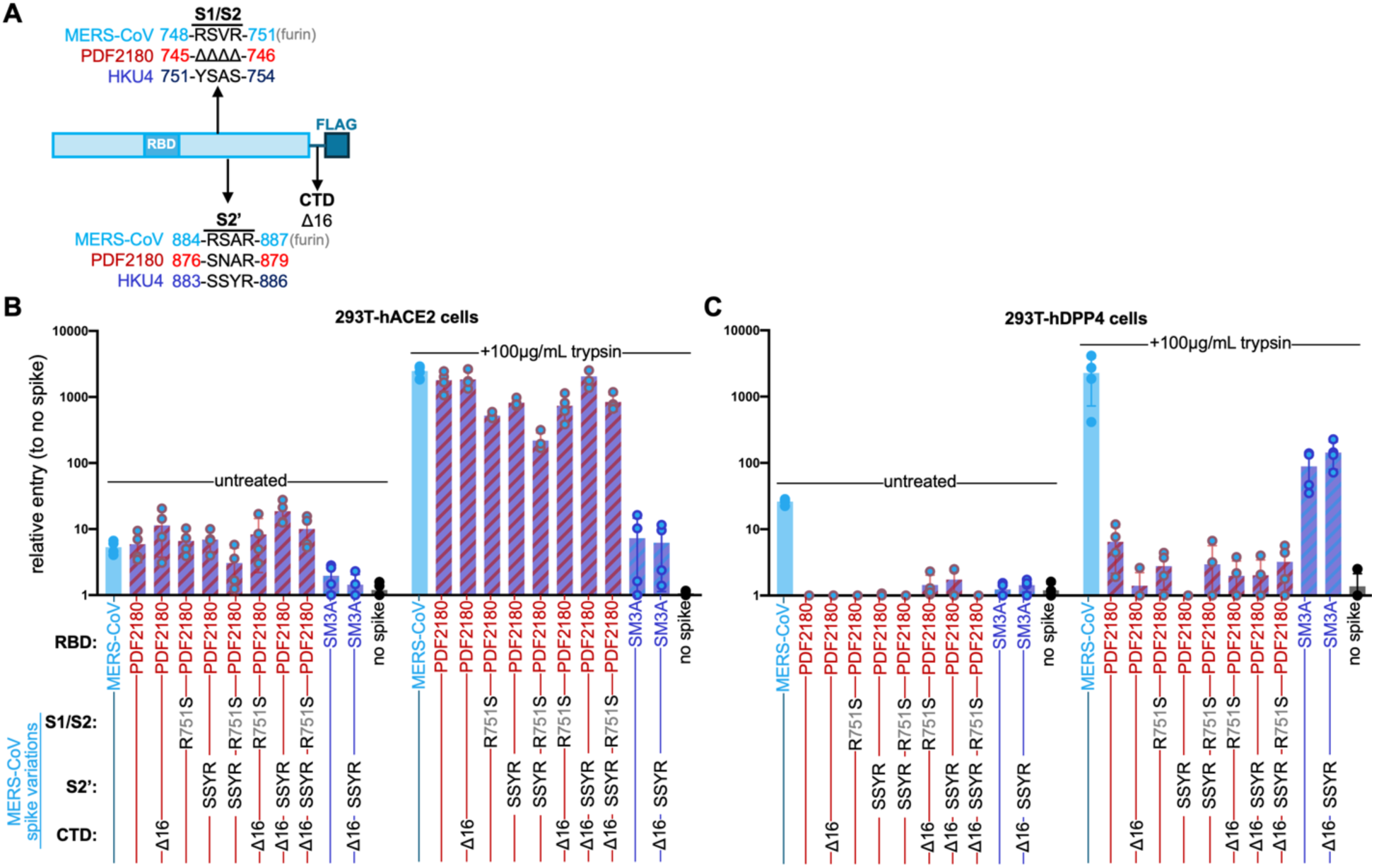
Optimizing MERS-CoV spike for MerbecoType. **(A)** Overview of S1/S2 and S2’ furin sites in MERS-CoV spike. **(B)** 293T cells transduced with human ACE2 or **(C)** human DPP4 were infected with indicated spikes and luciferase was measured for entry. Cells were infected in quadruplicate. For each graph, individual replicates are plotted as points, the mean is shown as a bar and lines indicate standard deviation.

**Figure S3.**
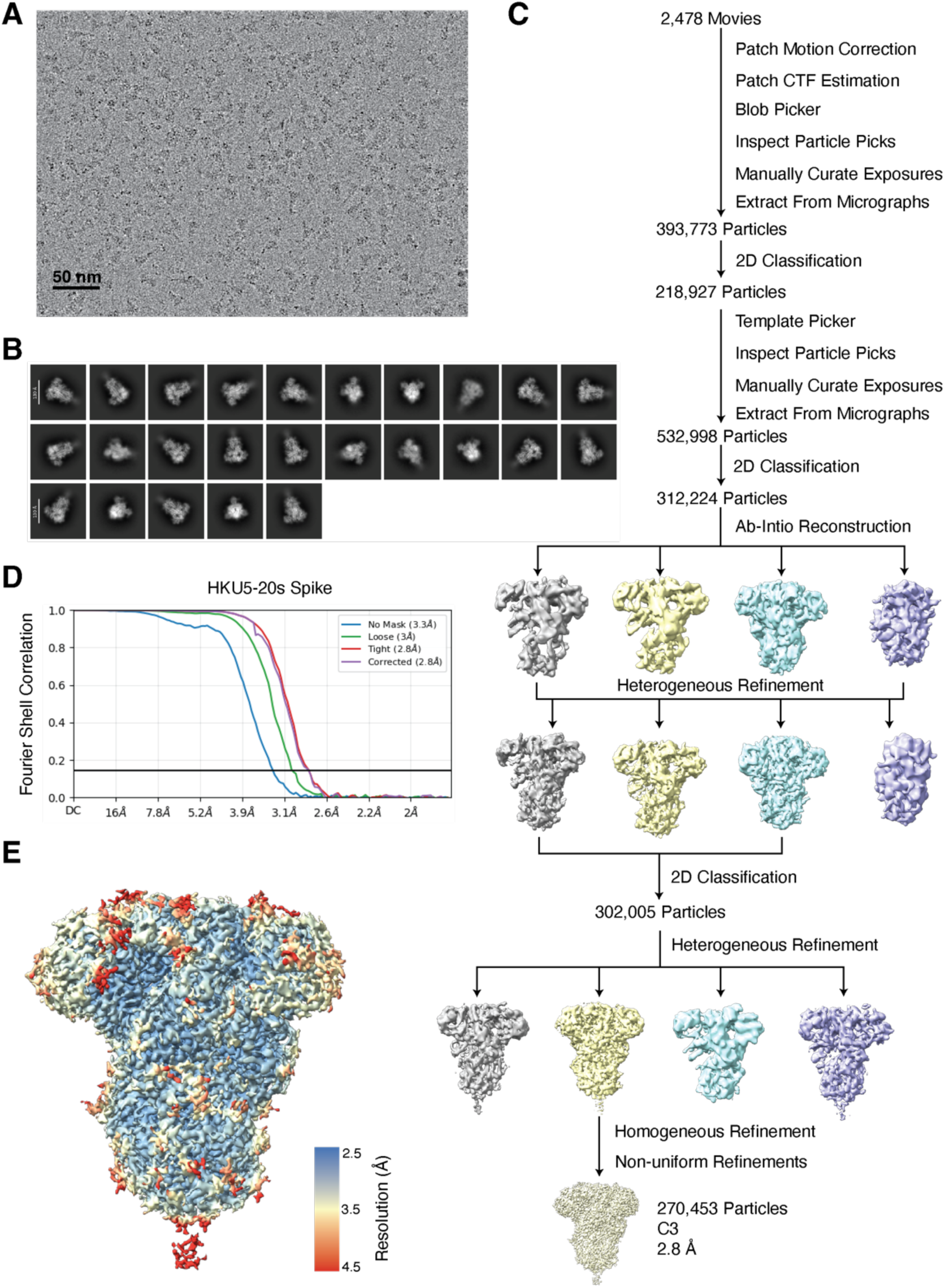
Cryo-EM workflow. **(A)** Representative micrograph. **(B)** Selected 2D classes. **(C)** Workflow of data processing. **(D)** Fourier shell correlation (FSC) plot of the final reconstruction. (E) Final reconstruction of HKU5-20s spike, colored from blue to red for local resolutions from high to low, respectively.

**TABLE S1.** Cryo-EM statistics for HKU5-20s Spike structure.

| Data Collection and processing |  |
| --- | --- |
| Microscope | Talos Arctica |
| Camera | Gatan K3 |
| Magnification | 45,000x |
| Voltage (keV) | 200 |
| Exposure (e/Å<sup>2</sup>) | 60 |
| Pixel size (Å) | 0.869 |
| Defocus Range (µm) | -1 to -3 |
| Initial Particle Images (no.) | 532,998 |
| Final Particle Images (no.) | 270,453 |
| Symmetry Imposed | C3 |
| Map Resolution (Å) | 2.8 |
| FSC Threshold | 0.143 |
| Map Resolution Range (Å) | 2.8-3.3 |
| Refinement |  |
| Initial Model Used | PDB 6NB3 |
| Model Resolution (Å) | 3.5 |
| FSC Threshold | 0.143 |
| Model composition |  |
| non-hydrogen atoms | 28,440 |
| protein residues | 3,548 |
| ligands | 63 |
| Average B-factors (Å<sup>2</sup>) |  |
| protein | 60.51 |
| ligands | 96.61 |
| R.M.S. deviations |  |
| Bond length (Å) | 0.002 |
| Bond angles (°) | 0.526 |
| Validation |  |
| MolProbity score | 1.6 |
| Clashscore | 6.7 |
| Rotamer outliers (%) | 1.3 |
| Ramachandran plot |  |
| Ramachandran favored (%) | 97.4 |
| Ramachandran allowed (%) | 2.6 |
| Ramachandran outliers (%) | 0 |
| PDB ID | 9D4T |
| EMDB ID | EMD-46569 |

